# Evolutionary conserved sap peptides derived from xylem-specific peptide precursors in woody angiosperms

**DOI:** 10.1101/2024.10.08.617308

**Authors:** Chang-Hung Chen, Pin-Chien Liou, Chih-Ching Lin, Shang-Che Kuo, Chia-Chen Wu, Ying-Chung Jimmy Lin, Ying-Lan Chen

## Abstract

Peptides act as long-distance mobile signals, transported through vascular sap to coordinate complex developmental processes. Since the tissue-specificity of peptide precursor gene expression is critical in determining peptide signaling function, we integrated vascular sap peptidomes with tissue-level transcriptomes to investigate the roles of sap peptides in two economically important woody plants, *Populus trichocarpa* and *Eucalyptus grandis*. Xylem exhibited the highest ratio of tissue-specific sap peptide precursor genes. Most of the sap peptides derived from xylem-specific precursor genes of *P. trichocarpa* and *E. grandis* were highly conserved throughout woody species selected from different clades in angiosperms, including magnoliids, rosids and asterids in eudicots. To further explore the conservation of these peptides, we examined the sap peptidome of *Cinnamomum kanehirae* (camphor tree), from the ancient clade with three xylem cell types. Approximately 90% of the peptides from xylem-specific precursors that were conserved between *P. trichocarpa* and *E. grandis*, were also conserved in the vascular sap of *C. kanehirae*, demonstrating a remarkably high conservation of these peptides across woody angiosperms. Most of the sap peptides conserved in these three woody species are also highly conserved across land plants, suggesting that these peptides may contribute to plant terrestrialization. Within the sap peptides from xylem-specific precursor genes, a total of 10 peptides were identical across all three woody plants. This substantial enrichment of xylem-specific precursor-derived peptides, along with their high conservation, suggests that these long-distance mobile peptides play a crucial role in secondary xylem development.

**One sentence summary:** Integration of sap peptidomic and tissue-level transcriptomic data revealed highly conserved long-distance mobile peptides derived from xylem-specific precursors across woody angiosperms.

## Introduction

Cell differentiation and development in multicellular organisms involve intercellular signaling sensed by membrane-bound receptors to modulate transcriptional regulatory networks for downstream responses (Fukuda, 2004; Sarkar et al., 2007; Reece and Campbell, 2011; Pierre-Jerome et al., 2018). In plants, vascular development is a multicellular process under complex regulation, including bifacial cell proliferation of single-layer initial cells (stem cells) followed by cell differentiation outward into phloem and inward into xylem. Previous studies have extensively revealed a conserved transcriptional regulatory mechanism of vascular development, mediated by the *VNS* (*VND*-, *NST*/*SND*-, *SMB* (*SOMBRERO*)-related protein) family master regulators (Kubo et al., 2005; Zhong et al., 2006; Zhong et al., 2008; Yamaguchi et al., 2010; Zhao et al., 2010; Xu et al., 2014), in land plants, especially in woody angiosperms (Tung et al., 2023; Chen et al., 2024). The *VNS* family members with their splice variants cooperatively regulate vasculature formation and secondary cell wall deposition through transcriptional regulatory networks majorly composed of *MYB* families (Li et al., 2012; Lin et al., 2013; Lin et al., 2017; Chen et al., 2019; Yeh et al., 2019; Wang et al., 2020). Despite the comprehensive understanding of transcriptional regulations in vascular development, the upstream intercellular signaling mechanisms remained largely unknown.

Peptides are hormone-like signals that regulate diverse biological functions through intercellular communications in plants (Chen et al., 2014; Motomitsu et al., 2015; Fukuda and Ohashi-Ito, 2019; Hirakawa and Sawa, 2019; Chen et al., 2020; Kim et al., 2021). Signaling peptides can be categorized as short- or long-distance mobile peptides based on their transport distance. Short-distance mobile peptides communicate with neighboring cells through symplastic or apoplastic pathways within a tissue or between tissues (Mitchum et al., 2008; De Smet et al., 2009). Long-distance mobile peptides travel exclusively via vascular sap to different tissues or organs, coordinating physiological processes in the entire plant (Okamoto et al., 2015; Carella et al., 2016; Okamoto et al., 2016; Takahashi et al., 2019; Fukuda and Hardtke, 2020).

CLAVATA3 (CLV3)/EMBRYO SURROUNDING REGION (CLE) and EPIDERMAL PATTERNING FACTOR-LIKE (EPFL) peptides are the only two peptide families reported as short-distance mobile peptides to coordinate vascular development through the regulation of procambial/cambial cells proliferation and differentiation into xylem and phloem (Tameshige et al., 2017; Hirakawa and Sawa, 2019; Fukuda and Hardtke, 2020; Yuan and Wang, 2021). TRACHEARY ELEMENT DIFFERENTIATION INHIBITORY FACTOR (TDIF) derived from CLE41/44 precursors inhibits the differentiation of tracheary elements and xylem from procambial cells (Ito et al., 2006; Hirakawa et al., 2008). The role of CLE25, CLE26 and CLE45 are related to protophloem differentiation (Hazak et al., 2017; Anne et al., 2018; Ren et al., 2019). CLE9/CLE10 regulates the periclinal cell division of xylem precursor cells and stomatal lineage cell division (Qian et al., 2018). EPFL4/6 peptides secreted from endodermis were perceived by ERECTA receptor in phloem cells to promote growth of inflorescence stem elongation (Uchida et al., 2012; Tameshige et al., 2017). In contrast to current discoveries of the regulation by short-distance mobile peptides on vascular development, long-distance mobile peptides have never been reported to regulate vascular development (Fukuda and Hardtke, 2020).

Even peptides can migrate throughout the whole plants, many studies have shown that peptides could affect the development of their precursor expressing tissues or the neighbor cells (Table S1). Simply put, if a peptide is generated from a precursor gene expressed in xylem, then this peptide may regulate xylem development. Furthermore, if this precursor gene is specifically expressed in xylem, then the peptide may have a higher chance to carry out a critical function on xylem development. Such tissue-dependent regulations were found in different parts of plant development. In *Arabidopsis*, CLV3 precursor genes were specifically expressed in shoot apical meristem (SAM), and the treatment of synthetic mature CLV3 induced occasional severe SAM reduction (Kondo et al., 2006). The gene expression of RGF1 was limited to quiescent center and columella stem cells in root apical meristem (RAM), and a dose-dependent RAM size increase was observed under the treatment of RGF1 peptide above 0.1 nM (Matsuzaki et al., 2010). In poplar, *CLE20* precursor gene was specifically expressed in secondary xylem, and its overexpression transgenics exhibited smaller radial width of secondary xylem area and smaller cell sizes of libriform fibers and vessel elements (Zhu et al., 2020). Overexpression of *CLE41*, while its precursor gene specifically expressed in phloem, caused intercalated phloem and highly organized vasculature (Etchells and Turner, 2010; Etchells et al., 2015). Thus, the tissue-specificity of precursor gene expression can be used to indicate the potential functions of mobile peptides.

In this study, our previously reported peptidomic vascular sap datasets of three woody plants (Chen et al., 2024), *Populus trichocarpa*, *Eucalyptus grandis* and *Cinnamomum kanehirae*, respectively were applied to investigate their potential conserved roles through the tissue-specificity and sequence conservation analysis. For *P. trichocarpa* and *E. grandis*, the vascular sap peptidomic and tissue-level transcriptomic analyses were integrated to analyze the tissue-specificity of peptide precursor gene expression. The conserved sap peptides derived from xylem-specifically expressed precursor genes of *P. trichocarpa* and *E. grandis* were further studied by their conservation throughout woody species selected from different clades in angiosperms. The vascular sap peptides of *C. kanehirae* (Chen et al., 2024), the ancient woody species from magnoliid, were also further studied for sequence conservation across angiosperm. Our results revealed a group of conserved long-distance mobile peptides originated from xylem serving the signaling role in xylem development.

## Results

### Identification of vascular sap peptides originated from tissue-specific precursors in *P. trichocarpa* and *E. grandis*

To reveal comprehensive profiles of long-distance mobile peptides, we analyzed the previous liquid chromatography tandem mass spectrometry (LC-MS/MS)-based sap peptidomic results of *P. trichocarpa* and *E. grandis* to reveal the tissue-specificity of the sap peptide precursor genes (Fig. 1) (Chen et al., 2024). In *P. trichocarpa*, we previously identified 1997 sap peptides that were derived from 801 peptide precursor genes (Table S2; Data S1). Since the expression locations of peptide precursors are critical to the functional role of their derived peptides (Table S1), we performed transcriptomic analysis of four tissues, xylem, phloem, leaves and young shoots in *P. trichocarpa* to characterize the expression location of the peptide precursor genes and their tissue-specificity (Fig. 1, orange part; Data S2) (Shi et al., 2017). If a precursor gene is specifically expressed in xylem (fold change > 1.5 and FDR < 0.05), then we would call such gene as a xylem-specific gene. In the 801 precursor genes, 32% (Fig. 1, middle left, 20 + 5 + 4 + 3%) were characterized as tissue-specific genes, and surprisingly, 20% were xylem-specific genes (Fig. 1, middle left). The dominant high ratio of xylem-specific precursor genes suggested that the major role of vascular sap peptides may contribute to xylem development, as known as wood formation in woody plants.

**Fig. 1.**
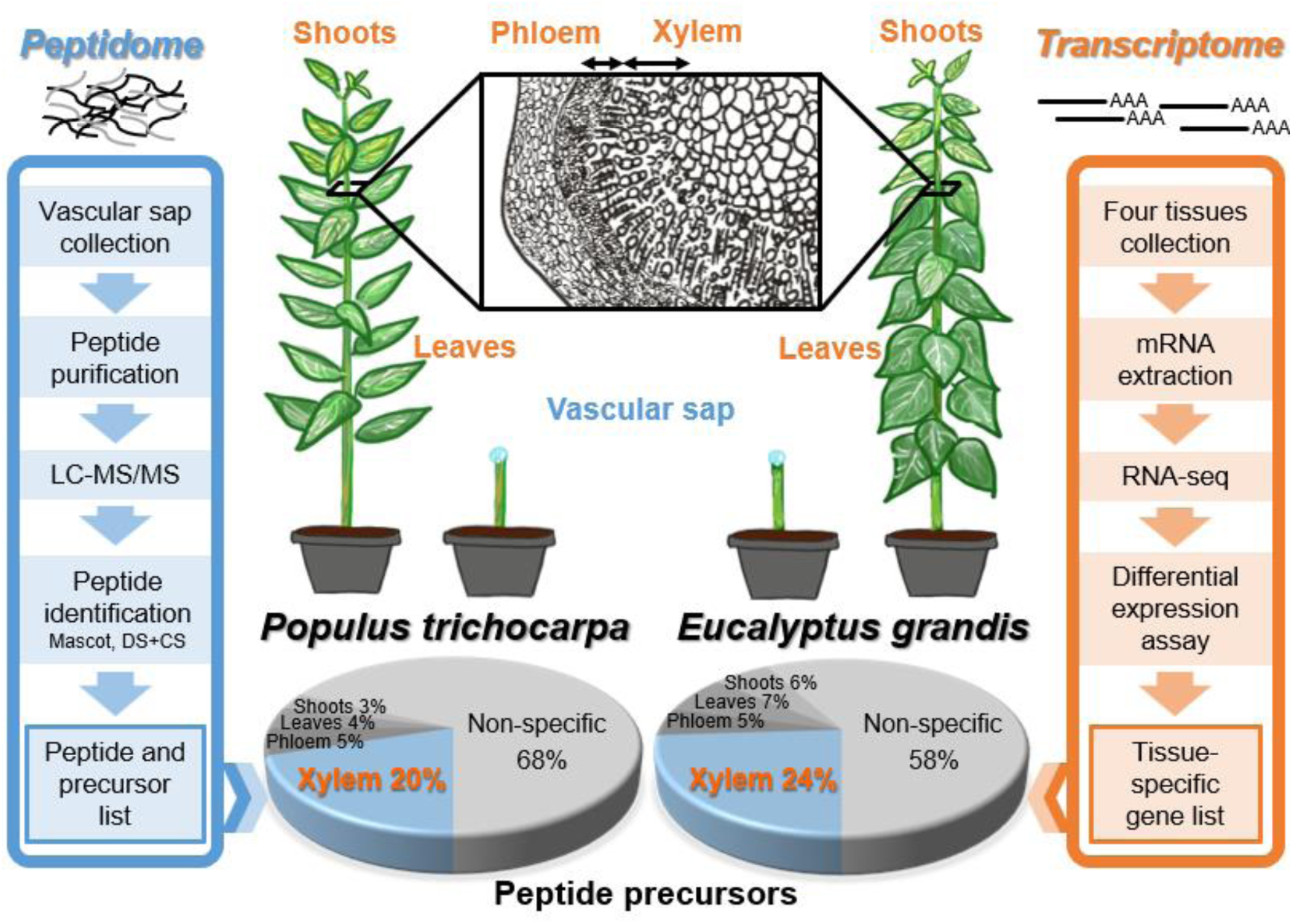
Integration of vascular sap peptidomic and tissue-level transcriptomic data to reveal the tissue-specificity of sap peptide precursor genes in *Populus trichocarpa* and *Eucalyptus grandis*. Workflows of tissue-level transcriptomic analyses and our previous vascular sap peptidomic analyses in *P. trichocarpa* and *E. grandis* (Chen et al., 2024). Transcripts extracted from four tissues (xylem, phloem, leaves, and young shoots) and vascular sap peptides previously collected from cutting stems were analyzed by RNA sequencing (RNA-seq) and liquid chromatography tandem mass spectrometry (LC-MS/MS), respectively. Previously, 801 and 229 vascular sap peptide precursors were identified in *P. trichocarpa* and *E. grandis*, respectively (Chen et al., 2024). The proportions of the sap peptide precursor genes specifically expressed in xylem, phloem, leaves and shoots were showed as 20% (164/801), 5% (40/801), 4% (33/801) and 3% (23/801) in *P. trichocarpa*, and 24% (56/229), 5% (11/229), 7% (15/229) and 6% (14/229) in *E. grandis*.

To investigate the evolutionary conservation of the high ratio of xylem-specific precursor genes, in addition to *P. trichocarpa* from malpighiales in eudicots, we also performed the four-tissue transcriptomic analysis in *E. grandis*, which is another eudicot woody species from myrtales (Fig. 1, middle right; Data S3). In *E. grandis*, 331 sap peptides were derived from 229 precursor genes (Table S2; Data S4). High ratio of xylem-specific precursor genes was also observed in *E. grandis* (Fig. 1, middle right). If xylem-specific genes are much more than other tissue-specific genes, then the xylem-specific precursor genes would tend to exhibit a higher ratio. In other words, the dominant ratio of xylem-specific precursor genes (Fig. 2A and C) may be caused by their original tissue-specificity ratio (Fig. 2B and D). We then normalized the precursor gene ratio by the percentage of tissue-specificity genes. Both *P. trichocarpa* and *E. grandis* showed significant tissue-specific enrichment ratio in xylem (Fig. 2E and F). In these two woody eudicots, the long-distance mobile peptides generated majorly from xylem-specific precursor genes, suggesting their evolutionary conserved functions in xylem development.

**Fig. 2.**
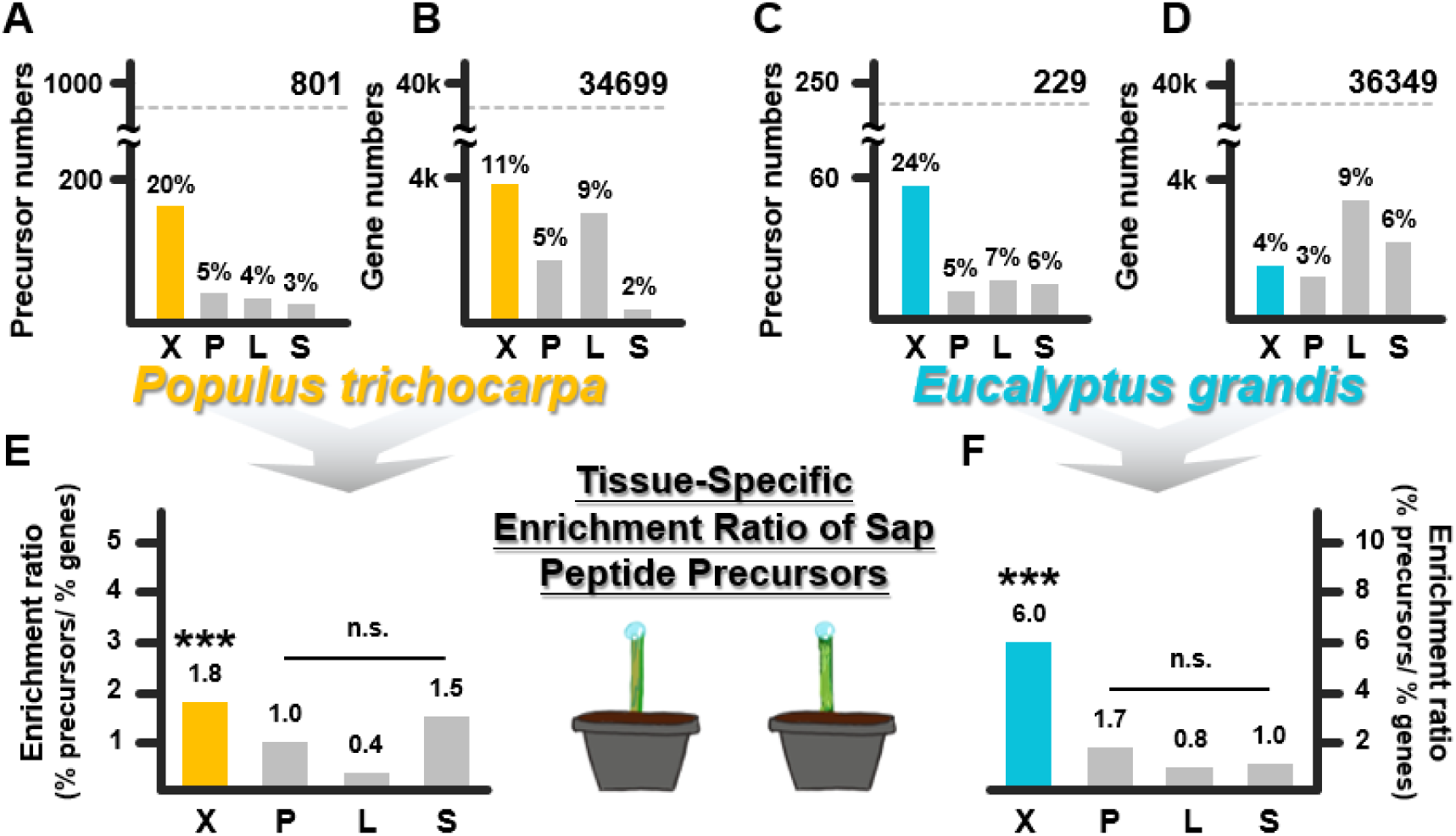
Enrichment analysis of vascular sap peptide precursors in tissue-specific genes of *Populus trichocarpa* and *Eucalyptus grandis*. The proportions of xylem-, phloem-, leaf- and shoot-specific gene numbers were represented as (**A**) 20%, 5%, 4% and 3% of total 801 vascular sap peptide precursors and (**B**) 11%, 5%, 9% and 2% of total 34,699 genes in *P. trichocarpa*, and (**C**) 24%, 5%, 7% and 6% of total 229 vascular sap peptide precursors and (**D**) 4%, 3%, 9% and 6% of total 36,349 genes in *E. grandis*. Tissue-specific enrichment ratios of xylem, phloem, leaves and shoots were, (**E**) 1.8, 1.0, 0.4 and 1.5 in *P. trichocarpa*, and (**F**) 6.0, 1.7, 0.8 and 1.0 in *E. grandis*, respectively. The enrichment ratios were calculated by the percentages of tissue-specific sap peptide precursors divided by the percentages of total tissue-specific genes. One-sided *p*-values calculated by normal approximation to the binomial test (proportions test) for the enrichment ratios of xylem, phloem, leaves and shoots were 2.9e-18 (***), 0.37, 0.99 and 0.24 in *P. trichocarpa*, and 2.9e-53 (***), 0.095, 0.92 and 0.49 in *E. grandis*, respectively. Three asterisks represent *p* < 0.001. X: xylem, P: phloem, L: leaves, S: young shoots.

Since the peptides were captured in vascular sap, specialized for long-distance migration, we examined the secretory signals on their precursors to find out whether these precursor proteins were secreted out of the cells to enter vascular sap. SignalP 5.0 were used for N-terminal signal peptide prediction (Almagro Armenteros et al., 2019), and 115 and 28 precursor proteins in *P. trichocarpa* and *E. grandis*, respectively, were predicted to contain secretory signals with high confidence, suggesting that these precursor proteins may be secreted into vascular sap to exert their functions (Data S1B, S4B).

### Exclusive conservation of sap peptides from xylem-specific precursors between *P. trichocarpa* and *E. grandis*

Besides the conserved high ratio of xylem-specific precursor genes, we then investigated the peptide sequence conservation between *P. trichocarpa* and *E. grandis*. We here consider the peptides as conserved if they share at least 70% sequence identity in two species. Total 246 and 111 conserved peptides were identified in *P. trichocarpa* and *E. grandis* (Fig. 3A; Table S2; Data S5). Among these conserved peptides, around 55-60% were from xylem-specific precursor genes (Fig. 3A), which is an even higher ratio comparing to the 20-24% dominant ratio of xylem-specific precursor genes (Fig. 2A and C). Such high proportion of xylem-specific precursor genes suggests that the majority of conserved sap peptides play a primary role in xylem development.

**Fig. 3.**
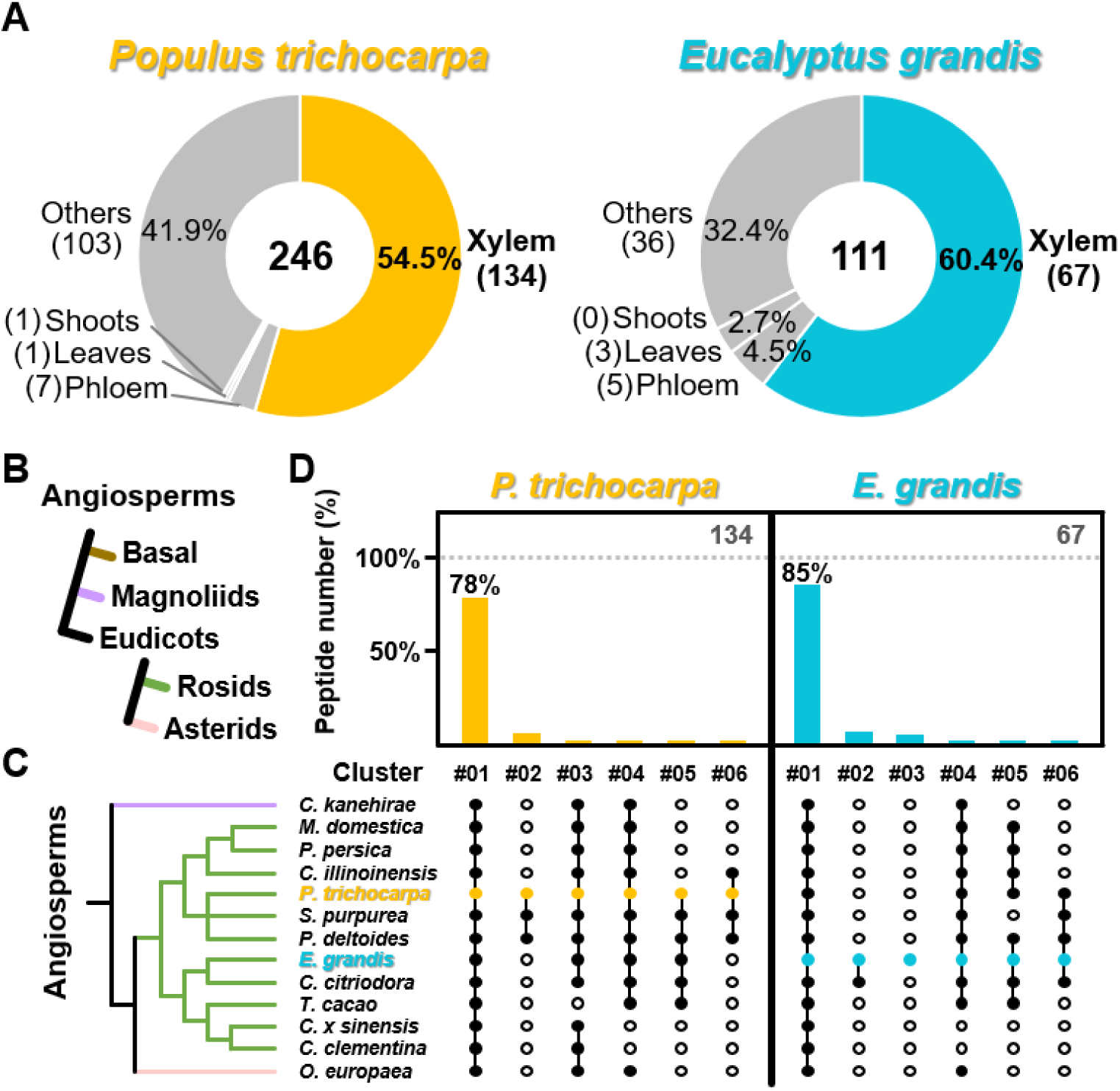
Conservation analysis of the conserved vascular sap peptides from xylem-specific precursor genes of *Populus trichocarpa* and *Eucalyptus grandis* in woody angiosperms. (**A**) Conservation analysis of identified sap peptides between *P. trichocarpa* and *E. grandis*. Total 246 and 111 conserved vascular sap peptides shared ≥ 70% sequence identity in *P. trichocarpa* and *E. grandis*, respectively. The proportions of conserved sap peptides from xylem-specific precursor genes were showed as 54.5% (134/246) and 60.4% (67/111) in *P. trichocarpa* and *E. grandis*, respectively. (**B**) Phylogenetic relationships of different clades including basal angiosperm, magnoliids, asterids and rosids of eudicots in angiosperms. (**C**) 13 woody species selected from different clades in angiosperms, *Cinnamomum kanehirae* from magnoliids, *Malus domestica*, *Prunus persica*, *Carya illinoinensis*, *P. trichocarpa*, *Salix purpurea, Populus deltoides WV94*, *E. grandis*, *Corymbia citriodora*, *Theobroma cacao*, *Citrus* × *sinensis* and *Citrus clementina* from rosids and *Olea europaea* from asterids. (**D**) UpSet plots of the Ptr-Egr conserved xylem-specific sap peptides containing the conserved peptide regions in 13 woody species. All peptide regions were generated from 134 or 67 conserved peptides of *P. trichocarpa* or *E. grandis*, respectively. Peptides with at least one peptide regions shared ≥ 90% sequence identity in other species were considered as conserved, indicated by filled circles. The proportions of *P. trichocarpa* or *E. grandis* sap peptides with peptide regions conserved in all 13 species were 78% (106/134) or 85% (57/67), respectively.

### Evolutionarily conserved sap peptides across woody angiosperms

Peptide production usually requires protease digestion. The protease recognition sites and cleavage sites locate near the peptide sequence with the length less than 10 residues (Farrokhi et al., 2008). Peptide sequences, protease recognition sequences and cleavage sites, here as “peptide regions”, are usually highly conserved for generating functional peptides (Farrokhi et al., 2008; Djordjevic et al., 2011; Katsir et al., 2011; Ni et al., 2011; Sharma et al., 2016). The analysis using peptide regions thus provides a better way to identify conserved precursors generating functional peptides. We further incorporated the 10 flanking residues on the C- and N-terminals next to the peptide sequences. A peptide region would be defined as conserved if it shares more than 90% alignment identity with other species, and we tested the evolutionary sequence conservation of the conserved peptides from xylem-specific precursor genes.

Xylem development in almost all angiosperm species involves the differentiation of three cell types as libriform fibers, vessel elements and ray parenchyma cells. Only very few species with reversal traits or in basal angiosperms possess tracheids and ray parenchyma cells (Cronk and Forest, 2017; Strijk et al., 2019). To study the peptide sequence conservation, we selected 13 woody species from angiosperms with the representing three xylem cell types. Eleven species, including *P. trichocarpa* and *E. grandis*, were from rosids in eudicots, and one species was from asterids in eudicots. Another species, *Cinnamomum kanehirae*, was selected from the most ancient clade with three xylem cell types, magnoliids (Fig. 3B and C). From the conserved peptide regions in *P. trichocarpa* and *E. grandis*, 78% (Fig. 3D, 106 of 134; Data S6A, S7A) and 85% (Fig. 3D, 57 of 67; Data S6B, S7B) Ptr-Egr-conserved peptides were found in all 13 species.

We then applied our previous sap peptidomic dataset of the camphor tree (*C. kanehirae*) to validate whether the conserved peptides could also be found in the vascular sap of the most ancient species among the 13 species (Chen et al., 2024). Total 899 *C. kanehirae* sap peptides were derived from 608 precursor genes (Data S8). Of the 106 (*P. trichocarpa*) and 57 (*E. grandis*) Ptr-Egr-conserved peptides, 100 and 51 peptides are also conserved in *C. kanehirae*, respectively, showing dramatically high ratio (∼90%, 100/106 and 51/57) of these peptides across angiosperms (Data S9). These 100 and 51 peptides meet the following five criteria (Fig. S1): (1) found in the vascular sap of *P. trichocarpa* or *E. grandis*; (2) conserved in the vascular sap of *E. grandis* and *P. trichocarpa*; (3) originated from xylem-specific precursor genes; (4) corresponding peptide region conserved in the 13 woody angiosperm species; and (5) conserved peptides found in the vascular sap of *C. kanehirae*. Our results suggest a group of conserved peptides across angiosperms, which may play as upstream signals to regulate xylem development.

### Peptide region conservation throughout land plants

Vascular system plays a critical role during land plant evolution for hundreds of million years (Fukuda and Ohashi-Ito, 2019; Lu et al., 2020; Ribeiro et al., 2020). With the specialized water-transporting and structural-supporting cells in xylem of vasculature, angiosperms are able to deliver essential nutrients and water over the whole plant body, which allows them to grow to large sizes with great heights. Since our results suggested that the conserved peptides in angiosperms may regulate xylem or vascular development, we next investigated the conservation of their corresponding peptide regions in other earlier vascular or non-vascular plants throughout the plant kingdom. Total 7 species were selected to represent (1) gymnosperms (*Gnetum montanum* and *Pinus taeda*) with different tracheary elements in vasculature, (2) spikemoss (*Selaginella moellendorffii*) as an ancient vascular plant species, (3) moss (*Physcomitrium patens*) and liverwort (*Marchantia polymorpha*) as early non-vascular land plants but still possessing primitive water-conducting and structural-supporting cells (Xu et al., 2014; Ohtani et al., 2017), and (4) algae (*Klebsormidium nitens* as a multicellular algae and *Chlamydomonas reinhardtii* as an unicellular algae) (Fig. 4A). We again used 90% sequence identity as the peptide region cutoff to examine the 100 and 51 Ptr-Egr-Cka conserved peptides in *P. trichocarpa* and *E. grandis*, respectively (Fig. 4B, S1). Total 78% (78/100 in *P. trichocarpa*) and 90% (46/51 in *E. grandis*) sap peptides have conserved peptide regions in seed plants (Fig. 4C; Data S10), and the peptides from both *P. trichocarpa* and *E. grandis* showed similar conservation ratios of 75% (75/100) and 71% (36/51), respectively, across vascular plants (Fig. 4D; Data S10). Around 55-70% of the Ptr-Egr-Cka conserved peptides still have conserved peptide regions in land plants (Fig. 4E; Data S10). A significant drop down to 16-21% appears upon the inclusion of algae (Fig. 4F; Data S10). These Ptr-Egr-Cka conserved peptides have conserved peptide regions within land plants with specialized water-conducting and structural-supporting cells, which suggested that these peptides may contribute to plant terrestrialization.

**Fig. 4.**
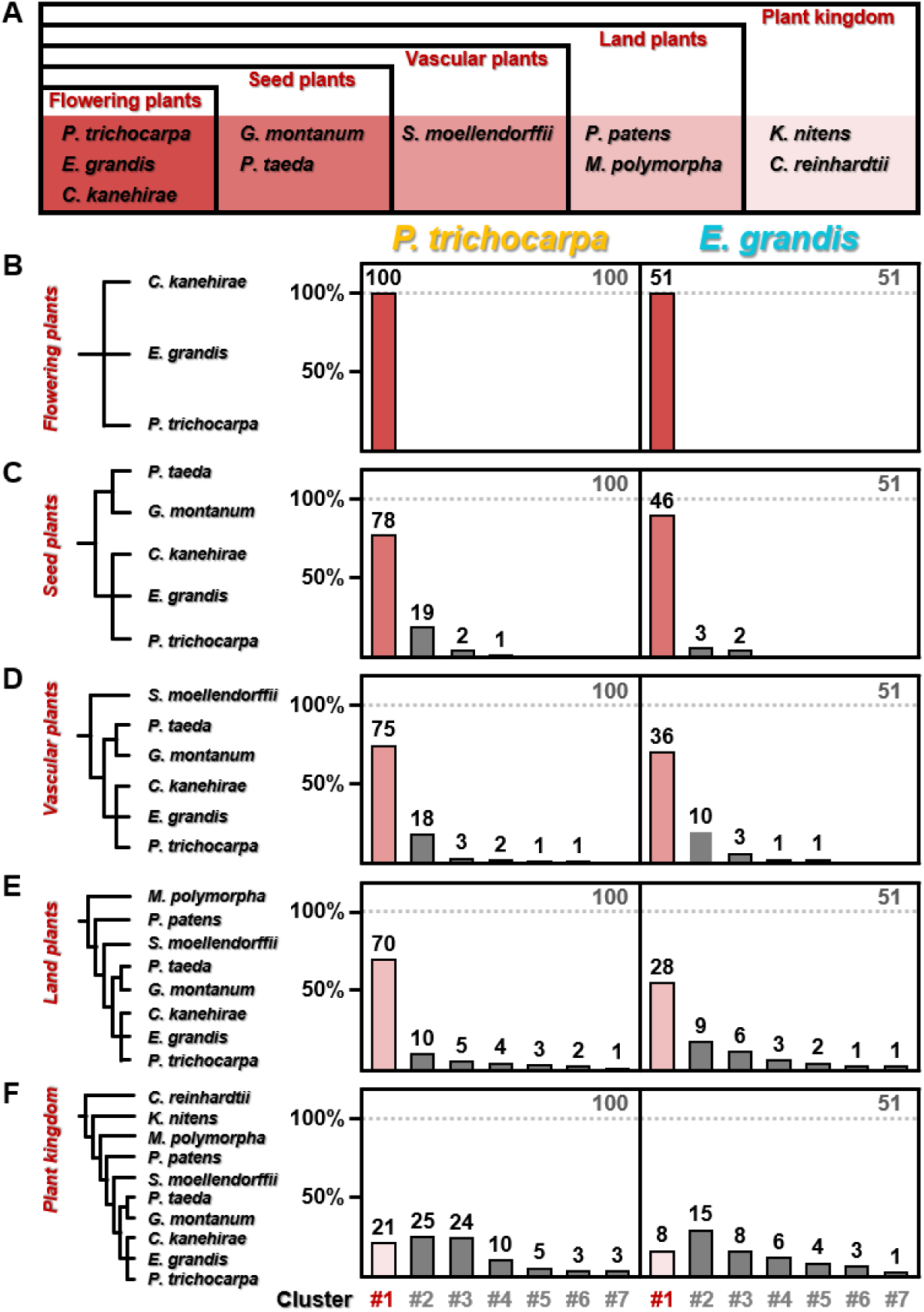
Conservation analysis of the Ptr-Egr-Cka conserved peptides across plant kingdom. (**A**) Species selected from the different clades of plant kingdom, including *Populus trichocarpa*, *Eucalyptus grandis*, *Cinnamomum kanehirae*, *Gnetum montanum*, *Pinus taeda*, *Selaginella moellendorffii*, *Physcomitrium patens*, *Marchantia polymorpha*, *Klebsormidium nitens* and *Chlamydomonas reinhardtii*. The peptide regions generated from Ptr-Egr-Cka conserved peptides were performed for the conservation analysis in (**B**) flowering plants (*P. trichocarpa*, *E. grandis* and *C. kanehirae*), (**C**) seed plants (*P. trichocarpa*, *E. grandis*, *C. kanehirae*, *G. montanum* and *P. taeda*), (**D**) vascular plants (*P. trichocarpa*, *E. grandis*, *C. kanehirae*, *G. montanum*, *P. taeda* and *S. moellendorffii*), (**E**) land plants (*P. trichocarpa*, *E. grandis*, *C. kanehirae*, *G. montanum*, *P. taeda*, *S. moellendorffii*, *P. patens* and *M. polymorpha*) and (**F**) plant kingdom (*P. trichocarpa*, *E. grandis*, *C. kanehirae*, *G. montanum*, *P. taeda*, *S. moellendorffii*, *P. patens*, *M. polymorpha*, *K. nitens* and *C. reinhardtii*). Peptides with at least one peptide region that shared ≥ 90% sequence identity in other species were considered as conserved. Peptides in different clusters have different conservation patterns, peptides in cluster #1 were conserved in all species of the selected clades.

### Identical vascular sap peptides across angiosperms

In our earlier study, the only one sap peptide found to be identical across six different angiosperm species (maize, camphor tree, tomato, rose gum, soybean, and poplar), <u>a</u>ngiosperm <u>sap</u> <u>p</u>eptide (ASAP), was discovered to be a functional peptide that may regulate various biological processes (Chen et al., 2024). To identify potential functional peptides that may regulate xylem development, we here also analyzed the sap peptides with identical sequences in *P. trichocarpa* and *E. grandis*. We found 28 identical peptides, and these peptides can be further grouped into 16 peptide families (Fig. 5A; Table S2; Data S11). Of these 28 identical peptides, 17 of them were from xylem-specific precursor genes in both *P. trichocarpa* and *E. grandis*, and 10 of them also have identical sequences in *C. kanehirae*, which can be grouped as 4 peptide families (Fig. 5B; Table S2; Data S11). Position weight matrix analyses revealed that the corresponding peptide regions of these 10 peptides, including ASAP, were highly conserved across the 13 woody angiosperm species (Fig. 6). We showed that ASAP functions as a long-distance mobile peptide, and xylem is one of its target destinations for ASAP (Chen et al., 2024). Based on the tissue-level transcriptomic analyses, we further confirmed that the precursor genes of ASAP are specifically expressed in xylem (Fig. S2A). Additionally, we used quantitative proteomic analysis across five different tissues revealed that the proteomic tissue-specificity of the ASAP precursors is consistent to the transcriptomic specificity (Fig. S2B). ASAP was shown to target to xylem and participate in the regulation of lignin and monolignol biosynthesis (Chen et al., 2024), the significant enrichment of the ASAP precursors in xylem further supports the idea that the expressing tissue of precursor gene is tightly correlated to the functional role of its derived peptides.

**Fig. 5.**
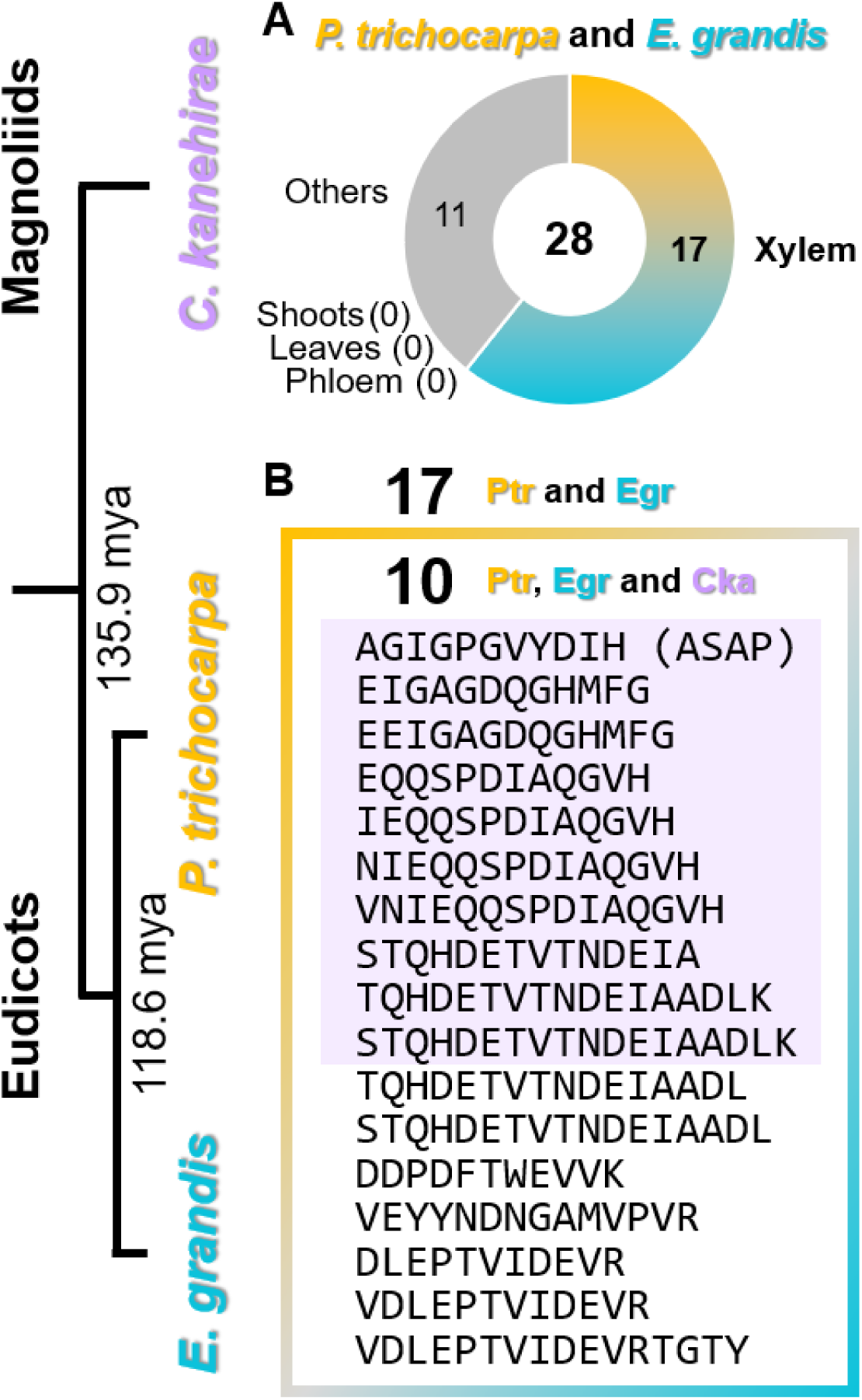
Identical vascular sap peptides in *Populus trichocarpa*, *Eucalyptus grandis* and *Cinnamomum kanehirae*. (**A**) Numbers of *P. trichocarpa* and *E. grandis* identical sap peptides and the tissue-specificity of their precursor genes. Total 28 peptides were identical in *P. trichocarpa* and *E. grandis*, including 17 peptides from xylem-specific precursor genes and 11 peptides from non-tissue-specific precursor genes. (**B**) Sequences of 17 identical peptides from xylem-specific precursor genes in *P. trichocarpa* and *E. grandis*, including 10 peptides that were also identical in *C. kanehirae* were listed. Left part of the figure shows the phylogenetic history of *P. trichocarpa* and *E. grandis* in rosids of eudicots and *C. kanehirae* from magnoliids (Magallón et al., 2015). ASAP, angiosperm sap peptide.

**Fig. 6.**
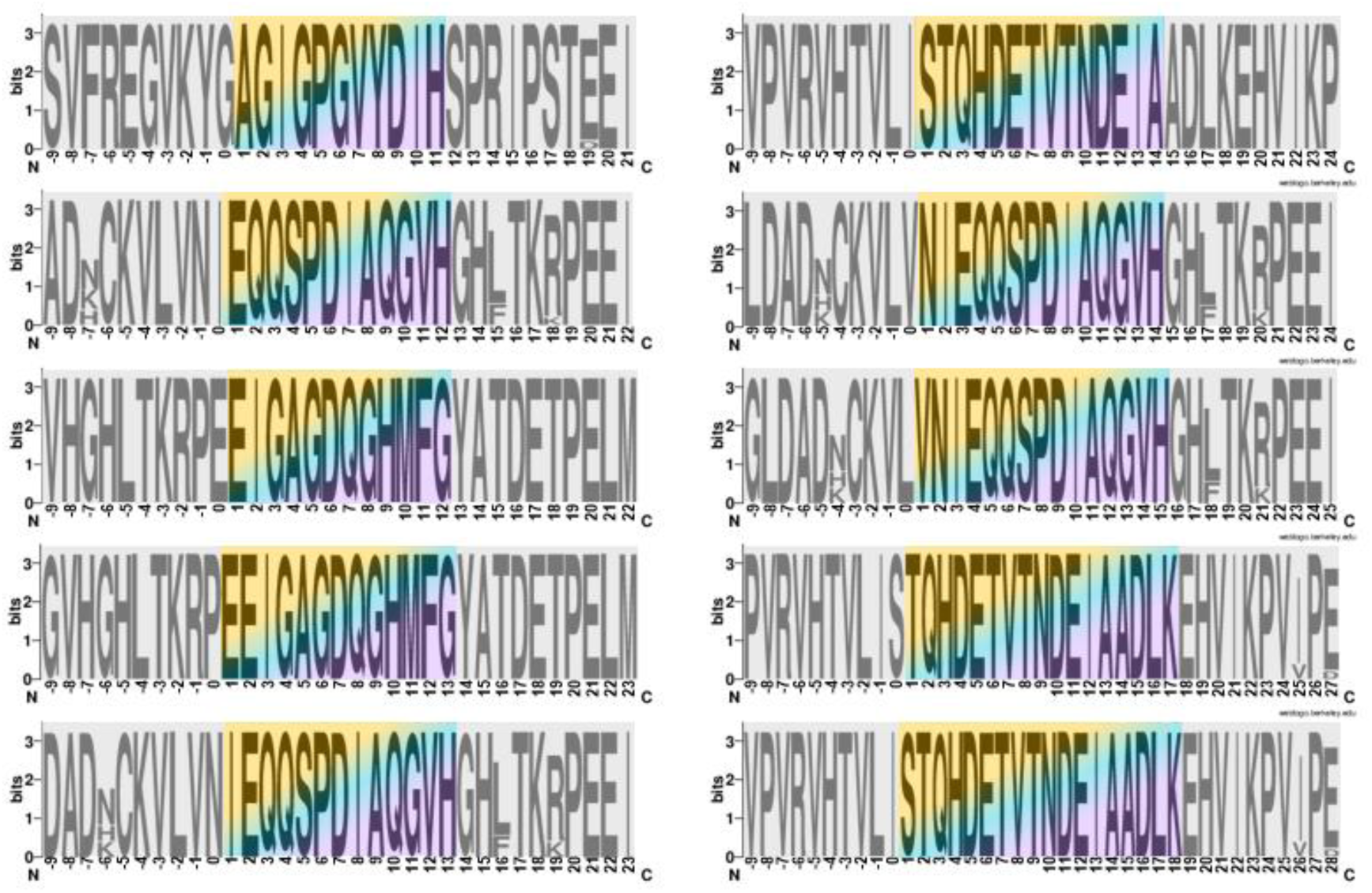
Position weight matrix (PWM) of the peptide regions of the Ptr-Egr-Cka identical peptides in woody angiosperms. PWMs of the 10 identical peptides with their 10 flanking residues in woody angiosperms. The BLASTP best hits of the conserved peptide regions in the 13 woody angiosperms (*C. kanehirae*, *M. domestica*, *P. persica*, *C. illinoinensis*, *P. trichocarpa*, *S. purpurea, P. deltoides WV94*, *E. grandis*, *C. citriodora*, *T. cacao*, *Citrus* × *sinensis*, *C. clementina* and *O. europaea*) were used for generating PWM by WebLogo (version 2.8.2).

## Discussion

Peptides transmit cell-to-cell communication to regulate many developmental processes in plants. Most of the previous studies focused on short-distance mobile peptides. For example, stomagen controls stomatal density (Sugano et al., 2010), RALF and LURE mediate pollen tube integrity and elongation (Okuda et al., 2009; Okuda et al., 2013; Ge et al., 2017), IDA governs floral organ abscission (Butenko et al., 2003), and CLE (TDIF) and EPFL regulate vascular development (Ito et al., 2006; Tameshige et al., 2017). Although sap peptides are primarily characterized by their mobility, previous studies have demonstrated that peptides can specifically regulate the development of their original tissues, where their precursor genes are expressed. Through the integration of sap peptide profiles and multi-tissue-level transcriptomes, we revealed hundreds of conserved long-distance mobile peptides generated majorly from xylem-specific genes of different families, which may contribute to the regulation of xylem development. Various types of peptide families were known to modulate the same developmental process. Short-distance mobile peptides CLE (TDIF) and EPFL are responsible for vascular patterning and differentiation (Ito et al., 2006; Tameshige et al., 2017). Four peptide families, RALF, STIG, PSK and LURE, regulate pollen tube growth (Tang et al., 2004; Okuda et al., 2009; Okuda et al., 2013; Stührwohldt et al., 2015; Ge et al., 2017). Treatment of synthetic CEP or PSK both increased root nodule numbers (Mohd-Radzman et al., 2016; Yu et al., 2022). EPF1, EPFL9 and SDD1 control stomatal development (Von Groll et al., 2002; Hunt et al., 2010). Thus, the different peptide families identified in vascular sap may work synergistically or antagonistically to control a same biological process, xylem development.

To exert long-distance migration, the peptide precursors are expected to be transported into vascular sap through protein secretory pathways (Alvarez et al., 2006; Ligat et al., 2011; Luo and Zhang, 2019). Most peptide precursors contain N-terminal secretory signals, and then are processed into peptides followed by secretion to the extracellular regions (Matsubayashi, 2011; Zhang et al., 2020). Only few wound-induced peptides are generated from non-secretory precursors and released to the extracellular region upon cell damage (Narvaez-Vasquez and Ryan, 2004; Hander et al., 2019; Chen et al., 2020). In contrast to past studies, further analyses based on our previous peptidomic profiles showed that 80∼90% of the sap peptide precursors, including all ASAP precursors in the three woody species, do not possess secretory signals (Data S1B, S4B). These non-secretory peptide precursors may be secreted by unconventional pathways (Chung and Zeng, 2017; van de Meene et al., 2017), or their peptides were released from broken cells (Hander et al., 2019; Hou et al., 2021). Xylem maturation process is accompanied by programmed cell death (PCD) (Bollhöner et al., 2018; Buono et al., 2019), suggesting that the vascular sap peptides originated from non-secretory xylem-specific precursors could be released during developmental PCD (Dafoe and Constabel, 2009).

Proteases, XYLEM CYSTEINE PEPTIDASE 1 (XCP1) and METACASPASE-9 (MC9), participated in xylo-developmental PCD and are regulated by the master regulators VND6 and VND7 of vascular development (Funk et al., 2002; Yamaguchi et al., 2011). XCP1 and MC9 were found to respectively produce the immunoregulatory peptide CAPE9 and oxidative stress induced peptide GRI in *Arabidopsis* (Wrzaczek et al., 2015; Chen et al., 2023), suggesting that XCP1 and MC9 may also execute peptide production during PCD in xylem development. The sequences of PCD-related proteases have been reported to be evolutionarily conserved (Coll et al., 2010), which suggests their cleavage sites and peptide products are conserved. We also found many highly conserved sap peptides and their flanking residues, the potential cleavage sites, in three woody angiosperms. Our results indicate that these PCD-related proteases may serve as the keys to recognize and process the evolutionarily conserved xylem-specific peptide precursors to generate upstream peptide signals for intra/inter-tissue communications.

Biological events in multicellular organisms are basically governed by two major steps, the intercellular signaling and the intracellular transcriptional regulation. In the past few decades, the intracellular transcriptional regulations of vascular development have been studied extensively, and the conserved transcription factor family *VNS* have been identified as master regulators in both moss and angiosperms (Xu et al., 2014; Nakano et al., 2015), demonstrating conserved regulatory mechanism in transcriptional regulation. In addition, the intercellular signaling molecules that regulate vascular development can be delivered over both short and long distances, such as hormones and peptides. Many members in CLE family are short-distance mobile peptides to control procambial/cambial cells proliferation and differentiation into xylem and phloem. Similar to *VNS* family, CLE family was also found to exert conserved functions in moss and angiosperms (Goad et al., 2017; Nemec-Venza et al., 2022). In addition, the co-receptor of CLE44 can interact with XVP to promote xylem differentiation through *VNS* family (Yang et al., 2020). However, a conserved intercellular and intracellular regulatory package for vascular development was known to be mostly mediated by only short-distance mobile peptides generated from neighboring cells. In this study, we uncovered a group of conserved long-distance mobile vascular sap peptides derived from stem-differentiating xylem-specific precursors that may significantly influence vascular development. Notably, these peptides are highly conserved across land plants, indicating their potential contribution to the process of plant terrestrialization. Our finding highlights the importance of understanding these peptides in unraveling the evolutionary mechanisms underlying vascular plant adaptation.

## Materials and Methods

### Plant materials and growth condition

Two-year-old *E. grandis* plants used for tissue-level transcriptomic analyses were grown from branch cuttings and maintained in a glasshouse at National Chung Hsing University. Four tissues, leaves, young shoots, phloem and xylem, were collected. Eight to nine-month-old *P. trichocarpa* and *E. grandis* plants used for detection of tissue-specific ASAP precursor proteins by proteomics analyses were grown from branch cuttings and maintained in the same glasshouse mentioned above in National Taiwan University. Five tissues, leaves, young shoots, roots, phloem and xylem, were collected. The collected leaves were fully expanded for both RNA and protein extraction. The young shoots were harvest as the first to third internodes and the first to eighth internodes for RNA extraction and protein extraction, respectively. The roots were harvested and washed cleanly for protein extraction. The whole bark scraped from the stem was used as phloem for RNA extraction and protein extraction. The stem-differentiating xylem was collected by scraping the surface of the debarked stems for both RNA and protein extraction.

### RNA extraction and sequencing

Four tissues (xylem, phloem, leaves and young shoots) of *E. grandis* were used for RNA extraction by CTAB extraction method. The CTAB extraction buffer was modified from Chang et al. (Chang et al., 1993), containing 2% (w/v) hexadecyltrimethylammonium bromide, 0.1 M Tris-HCl (pH 9.0), 25 mM EDTA (pH 8.0), 2 M NaCl and 1% (w/v) PVP-40. Two percent (v/v) of β-mercaptoethanol and 50 mM ascorbic acid was added to the extraction buffer before use. Each tissue sample was ground into fine powder in liquid nitrogen, and 0.5 g powder of each sample was stored in 1.5-ml tubes. The 5-ml pre-warmed (65℃) CTAB extraction buffer was used to resuspend the power of each sample followed by vortexing to fully mix the powder and the buffer with 65℃ incubation for 10 min. After centrifugation at 12,000 ×g for 5 min at room temperature, the supernatant was transferred to new 1.5-ml tubes, and an equal volume of chloroform: isoamyl alcohol (24:1) was added. Each sample was vortexed and then centrifuged for 10 min at 12,000 ×g at room temperature. The chloroform extraction step was repeated twice. The aqueous phase was mixed with one-third volume of 8 M lithium chloride and kept at 4℃ overnight. The RNA pellet was collected by centrifuging for 20 min at 12000 ×g at 4℃. The RNA in the pellet was further purified by the Qiagen RNeasy Plant RNA isolation kit. The RNA library construction was performed using NEBNext^®^ Ultra™ II Directional RNA Library Prep Kit for Illumina^®^ followed by RNA sequencing using Illumina HiSeq X Ten.

### Differentially expressed gene analysis

RNA-seq datasets of xylem, phloem, leaves and young shoots from *P. trichocarpa* and *E. grandis* were used to identify the tissue-specifically expressed genes. The dataset of *P. trichocarpa* was downloaded from NCBI database (GEO accession number: GSE81077) (Shi et al., 2017). The dataset of *E. grandis* was generated in this study. The RNA-seq raw reads were processed by fastp to trim the adapters and to remove low-quality reads containing more than 40% of bases with Phred quality score below 15 (Chen et al., 2018), resulting in clean reads to obtain transcripts per million (TPM) values by transcript abundance quantification using RSEM (Li and Dewey, 2011). FDR was calculated based on unnormalized gene read counts of the RSEM output through DESeq2 package in R (Love et al., 2014), and the genes with read counts less than 10 were removed from DEG candidates. A gene is defined as a xylem-specifically expressed gene if this gene meets the following two criteria: all three FDR values (xylem vs phloem; xylem vs leaves; xylem vs young shoots) were lower than 0.05, and its transcript abundance is at least 1.5 times higher than in the other three tissues (Lin et al., 2017). Further, the tissue-specificity of the identified sap peptides are defined by the tissue-specificity of their precursor genes. For example, if a peptide has only one precursor gene that was specifically expressed in xylem, this peptide would be defined as from xylem-specific precursor. If a peptide has multiple precursor genes with at least one precursor is xylem-specifically expressed gene while others have no tissue-specificity, this peptide would also be defined as xylem-specific. If a peptide has multiple precursors with different tissue-specificity, then this peptide would be defined as non-specific.

### Peptide conservation analysis

Conservation analysis of the sap peptides between *P. trichocarpa* and *E. grandis* was performed by BLASTP using default settings and e-value < 100 (Altschul et al., 1997). Peptides would be considered as conserved if their sequence identity were equal to or higher than 70% in *P. trichocarpa* and *E. grandis* under full-length alignment. The same method was used for analyzing the sap peptide conservation in *C. kanehirae*.

### Peptide region conservation analysis

Peptide regions were defined as the combined sequences of peptide sequences and the 10 flanking residues on the C- and N-terminals. The peptide regions of *P. trichocarpa* and *E. grandis* were respectively used to search against the protein databases of different species by BLASTP with the same parameters used in Phytozome (e-value < 0.001, BLOSUM45 substitution matrix and default settings) (Goodstein et al., 2012). A peptide would be considered as conserved in a species if at least one of its peptide regions have BLASTP hits with at least 90% sequence identity, and that the alignment region includes the entire peptide sequence.

The protein fasta files of each species were downloaded from Phytozome (*Cinnamomum kanehirae* v3, *Malus domestica* v1.1, *Prunus persica* v2.1, *Carya illinoinensis* v1.1, *Populus trichocarpa* v4.1, *Salix purpurea* v5.1, *Populus deltoides WV94* v2.1, *Eucalyptus grandis* v2.0, *Corymbia citriodora* v2.1, *Theobroma cacao* v2.1, *Citrus* × *sinensis* v1.1, *Citrus clementina* v1.0, *Olea europaea* v1.0, *Selaginella moellendorffii* v1.0, *Physcomitrium patens* v3.3, *Marchantia polymorpha* v3.1 and *Chlamydomonas reinhardtii* v5.6) (Goodstein et al., 2012), TreeGenes (*Gnetum montanum* v1.0 and *Pinus taeda* v2.01) or NCBI (*Klebsormidium nitens*, GenBank accession: GCA_000708835.1).

### Position weight matrix of conserved peptide regions

Peptide regions of the 10 identical sap peptides of *P. trichocarpa*, *E. grandis* and *C. kanehirae* along with the homologous sequences with highest sequence similarity in total 13 woody angiosperms (*C. kanehirae*, *M. domestica*, *P. persica*, *C. illinoinensis*, *P. trichocarpa*, *S. purpurea, P. deltoides WV94*, *E. grandis*, *C. citriodora*, *T. cacao*, *C.* × *sinensis*, *C. clementina* and *O. europaea*) were used to generate the position weight matrices (PWMs) by the default settings of WebLogo (version 2.8.2) (Crooks et al., 2004).

### Protein extraction, digestion and tryptic peptides purification

Five tissues (xylem, phloem, leaves, roots and young shoots) of *P. trichocarpa* and *E. grandis* were used for protein extraction using a previous published method (Wang et al., 2006). Protein pellet was dissolved by a lysis buffer containing 5% SDS in 50 mM Tris-HCl (pH 8.0), and then the concentrations of dissolved proteins were measured using BCA protein assay (Thermo Fisher Scientific). Before trypsin digestion, the disulfide bonds of proteins were reduced with 10 mM tris (2-carboxyethyl) phosphine (TCEP) and alkylated with 40 mM chloroacetamide at 45 ℃ for 10 min. Each protein sample was digested into peptides by trypsin in a S-trap micro column to remove the incompletely digested proteins and trypsin. The tryptic peptides were acidified by trifluoroacetic acid and desalted by SDB-XC StageTips (catalog no. 2340; 3M). The desalted samples were dried by a vacuum centrifugation concentrator. The dried samples were dissolved in 0.1% formic acid (FA) and centrifuged at 13,000 ×g for 10 min at 4℃. The supernatant was transferred into a sample vial, and analyzed by LC-MS/MS analysis.

### LC-MS/MS-based proteomic analyses

Five tissues of digested peptide samples of *P. trichocarpa* were analyzed by a Dionex 3000 UPLC system coupled with a Q Exactive Hybrid Quadrupole-Orbitrap mass spectrometer (Thermo Fisher Scientific). Peptides were separated on a 25-cm PepMap C18 column packed with 2-μm particles (Thermo Fisher Scientific) using the mobile buffer consisted of 0.1% FA in ultra-pure water with an eluting buffer of 0.1% FA in 100% acetonitrile (ACN) with a linear 60 min gradient of 5 to 25% ACN/0.1% FA at a flow rate of 300 nl/min. The mass spectrometer was first operated in data-dependent acquisition (DDA) mode for spectral library construction, and then performed data-independent acquisition (DIA) mode for protein quantification. For DDA mode, a full scan (from m/z 350-1600 with the resolution of 70,000 at m/z 200) was followed by higher-energy collisional dissociation fragmentation using the 10 most intense ions with normalized collision energy of 27 and the resolution of 17,500 at m/z 200. The automatic gain control (AGC) value of MS was set as 1e5 with the max injection time as 120 ms. The isolation window was set as 2.0 while the dynamic exclusion duration as 20 s. For DIA mode, a full scan of mass range was set as m/z 350-950, and 28 windows of 20 m/z scanning from 400 to 900 m/z were used with an overlap of 1 Da. Resolution was set as 17,500, injection time as 50 ms, and the normalized collision energy was 27%.

Five tissues of digested peptides of *E. grandis* were analyzed by an Easy nLC 1200 coupled with Orbitrap Fusion Lumos mass spectrometer (Thermo Fisher Scientific). Peptides were separated by an Acclaim PepMap 100 C18 trap column (75 µm x 2.0 cm, 3 µm, 100 Å , Thermo Fisher Scientific) and an Acclaim PepMap RSLC C18 nano LC column (75 µm x 25 cm, 2 µm, 100 Å ) using the mobile buffer consisted of 0.1% FA in ultra-pure water with an eluting buffer of 0.1% FA in 100% ACN with 5 to 25% ACN/0.1% FA at a flow rate of 300 nL/min. The 60 min linear gradient was used.

For DDA mode, a full scan (m/z 350-1600 with the resolution of 120,000 at m/z 200) was followed by MS/MS fragmentation in a 3 sec cycle time. The maximum ion injection time was set 50 as ms. The MS/MS acquisitions were performed using 1.4 Da isolation window with 50,000 AGC value, 35% normalized collision energy (NCE), maximum injection time as 120 ms, and 15,000 resolving power. For DIA mode, a full scan (m/z 400-1000 with the resolution of 120,000 at m/z 200), and followed by 40 MS/MS scans. The MS scan was performed with 120,000 resolving power over the m/z range 350 to 1050 and dynamic exclusion enabled. The DIA MS/MS acquisitions were performed using 15 Da isolation window with an overlap of 1 Da, 27% NCE and 15,000 resolving power.

### Target-decoy database

Target-decoy database was used for MS/MS ion searching and false discovery rate (FDR) evaluation (Elias and Gygi, 2007), which was generated by combining the protein sequences with the randomized protein sequences (decoy) for each species using Trans Proteomics Pipeline (TPP) version 5.1 (Pedrioli, 2010). The protein sequences of *P. trichocarpa* and *E. grandis* (*Populus trichocarpa* v4.1; *Eucalyptus grandis* v2.0) were downloaded from Phytozome (Goodstein et al., 2012).

### MS data processing and database searching

All DIA raw files and spectral libraries were processed and generated by DIA-NN (version 1.8) (Demichev et al., 2020). Trypsin/P was set as a specific protease and allowed up to two missed cleavages. FDR at peptide and protein level was set to 1%. MS1 accuracy was set to 10 ppm, and mass accuracy was set to 20 ppm. Peptide length range was set to 7-30 amino acid residues. Precursor charge range was set as 2-4.

Precursor m/z range was set to 350-1050 m/z, and fragment ion m/z range was set to 200-1800. Neural network classifier was set as single-pass mode and quantification strategy was set as Robust LC. Match between run function was enabled, and RT-dependent mode was selected as cross-run normalization. The expression levels of ASAP precursor proteins in different tissues were extracted from the main output file of DIA-NN using the DiaNN R Package (version: diann_1.0.1).

## Acknowledgments

We thank Dr. Ying-Hsuan Sun for providing the *E. grandis* plant materials for RNA extraction, Cing Shih for mRNA extraction and library construction.

## Author contributions

Y.L.C., Y.J.L. supervised the project and designed experiments. C.H.C., S.C.K. performed the transcriptomic data analysis. Y.L.C., C.H.C. analyzed the peptidomic and proteomic datasets. Y.L.C., P.C.L. performed the proteomic experiments. C.H.C., P.C.L., Y.L.C., Y.J.L., C.C.L. constructed all figures. Y.L.C., Y.J.L., C.H.C., P.C.L., C.C.W. wrote the manuscript.

## Funding

Taiwan MOST Young Scholar Fellowship Einstein Program 109-2636-B-006-009 (Y.L.C.)

Taiwan MOST Young Scholar Fellowship Einstein Program 110-2636-B-006-008 (Y.L.C.)

Higher Education Sprout Project, Ministry of Education to the Headquarters of University Advancement at National Cheng Kung University (Y.L.C.)

Taiwan MOST International Young Scholar Fellowship Program 110-2628-B-002-026 (Y.J.L.)

Taiwan MOST International Young Scholar Fellowship Program 111-2628-B-002-020 (Y.J.L.)

## Data availability

Tissue-level RNA-seq raw reads were downloaded from or deposited at GEO of NCBI, accession number GSE81077 and GSE174856 for *P. trichocarpa* and *E. grandis*, respectively. All peptidomic data were downloaded from or deposited at the ProteomeXchange Consortium via the PRIDE partner repository with the dataset identifier PXD032088 (Perez-Riverol et al., 2022). All other data needed to evaluate the conclusions in the paper are present in the paper and/or the Supplementary Materials.

## Supplementary Figures

**Fig. S1.**
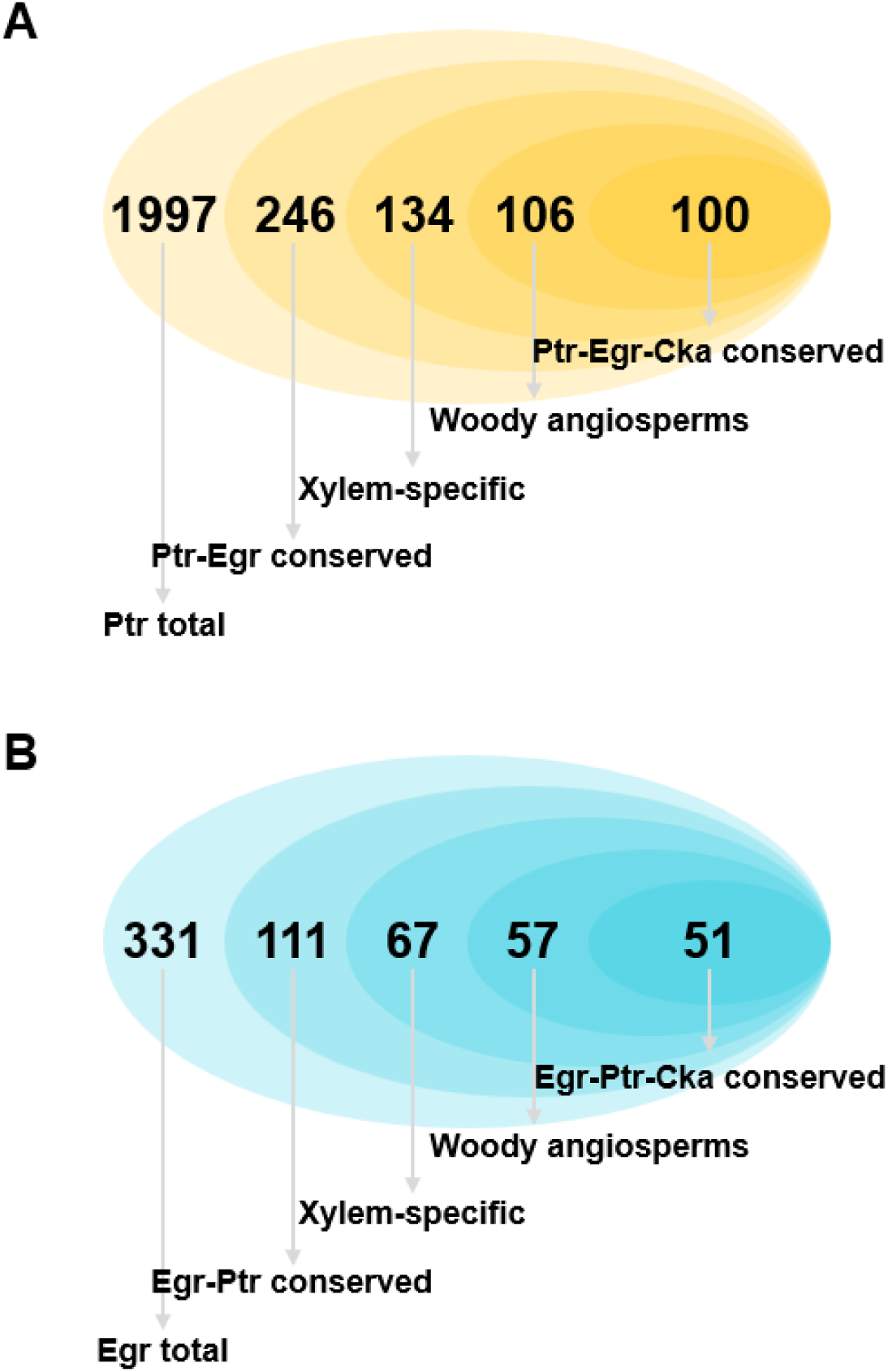
The number of previously identified vascular sap peptides in *Populus trichocarpa* and *Eucalyptus grandis* selected by different criteria. (**A**) In *P. trichocarpa*, total 1997 peptides were previously detected in the vascular sap (Chen et al., 2024), 246 of the 1997 peptides were conserved in *P. trichocarpa* and *E. grandis*, 134 of the 246 peptides were originated from xylem-specific precursor genes, 106 of the 134 peptides had at least one peptide regions conserved in all 13 woody angiosperms, and 100 of the 106 peptides were conserved in the vascular sap of *C. kanehirae*. (**B**) In *E. grandis*, total 331 peptides were previously detected in the vascular sap (Chen et al., 2024), 111 of the 331 peptides were conserved in *E. grandis* and *P. trichocarpa*, 67 of the 111 peptides were originated from xylem- specific precursor genes, 57 of the 67 peptides had at least one peptide regions conserved in all 13 woody angiosperms, and 51 of the 57 peptides were conserved in the vascular sap of *C. kanehirae*.

**Fig. S2.**
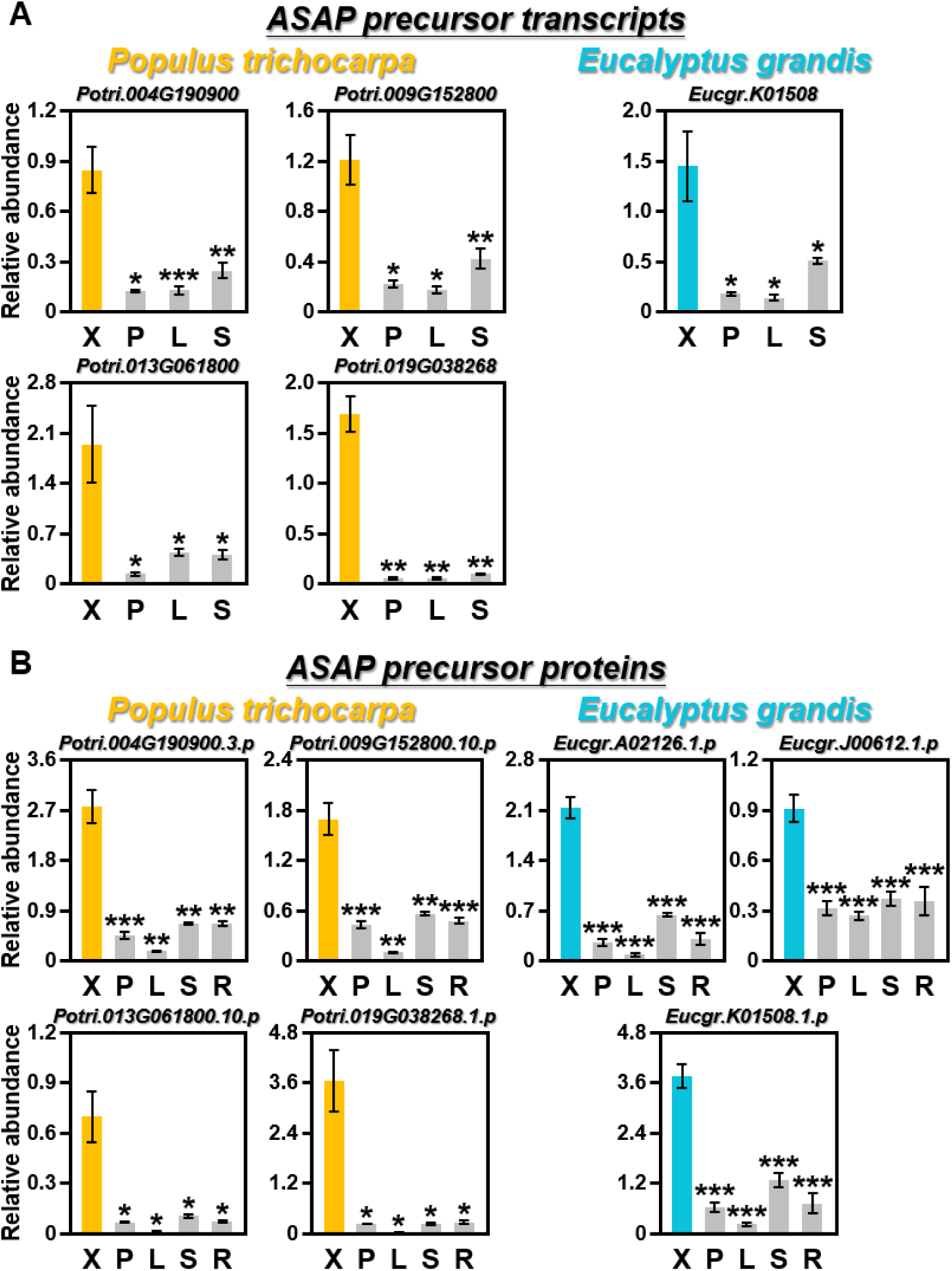
Tissue-level expression of ASAP precursor transcripts and proteins in *P. trichocarpa* and *E. grandis*. (**A**) Expression level of ASAP precursor transcripts in *P. trichocarpa* and *E. grandis* based on the four-tissue transcriptomic analyses. X: xylem, P: phloem, L: leaves, S: young shoots. One, two and three asterisks represent Student’s *t*-test *p* < 0.05, 0.01 and 0.001, respectively. (**B**) Expression level of ASAP precursor proteins in *P. trichocarpa* and *E. grandis*. Quantitative proteomic analyses of five tissues, xylem (X), phloem (P), leaves (L), young shoots (S) and roots (R) of *P. trichocarpa* and *E. grandis* were performed by LC-MS/MS with a data-independent acquisition (DIA) method. The relative abundances of all ASAP precursor proteins in five tissues were calculated from the raw data using DIA- NN (version 1.8). Three and four biological replicates were performed for *P. trichocarpa* and *E. grandis* plants, respectively. One, two and three asterisks represent Student’s *t*-test *p* < 0.05, 0.01 and 0.001, respectively. ASAP, angiosperm sap peptide.

## Supplementary Tables

**Table S1-1.** Expressing and activating tissues of peptides during plant development.

| Peptide Name | Expressing tissue | Activating tissue | Function related to | Species | Ref. (PMID) |
| --- | --- | --- | --- | --- | --- |
| CEP3 | Roots | Roots | Lateral root development | *A. thaliana* | 29366684<br>24179096 |
| CLV3 | SAM* | SAM* | SAM* development | *A. thaliana* | 16902141 |
| CLE5/6 | Leaves<br>Flowers | Leaves | Leaf development | *A. thaliana* | 31245702 |
| CLE8 | Embryos<br>Endosperm | Embryos<br>Endosperm | Embryo/endosperm development | *A. thaliana* | 22427333 |
| CLE9/10 | Leaf stomata | Leaf stomata | Stomatal development | *A. thaliana* | 30518839 |
| CLE10 | Root vascular | Root vascular | Root vascular development | *A. thaliana* | 20802224 |
| CLE12/13 | Root nodules | Root nodules | Root nodule development | *M. truncatula* | 20348212 |
| CLE14/20 | Roots | Roots | RAM* development | *A. thaliana* | 20697738 |
| CLE20 | Xylem | Xylem<br>Phloem<br>Cambium | Vascular development | *P. trichocarpa* | 31199056 |
| CLE25 | Phloem<br>Procambium | Phloem | Phloem development | *A. thaliana* | 31127689 |
| CLE26/45 | RAM*<br>Root phloem | Root phloem | Root phloem development | *A. thaliana* | 29789310<br>31327718 |
| CLE41/44 | Phloem | Procambium | Vascular development | *A. thaliana* | 16902140<br>18812507<br>26414535 |
| CIF1/2 | Root stele | Root stele | Root Casparian strip formation | *A. thaliana* | 28104889 |
| EPF2 | Leaf stomata | Leaf stomata | Stomata density on leaf | *A. thaliana* | 19398336 |

\* SAM: Shoot apical meristem; RAM: Root apical meristem

**Table S1-2.** Expressing and activating tissues of peptides during plant development.

| Peptide Name | Expressing tissue | Activating tissue | Function related to | Species | Ref. (PMID) |
| --- | --- | --- | --- | --- | --- |
| EPFL4/6 | Endodermis | Phloem | Inflorescence stem development | *A. thaliana* | 27965367<br>22474391 |
| ESF1 | ESR* | ESR* | Embryo development | *A. thaliana* | 26195729<br>24723605 |
| FCP1 | SAM*<br>RAM* | SAM*<br>RAM* | SAM* and RAM* maintenance | *O. sativa* | 18676878 |
| GAD1 | Panicles | Panicles | Panicle development | *O. sativa* | 27634315 |
| GLV6 | RAM*<br>LRP* | LRP* | Lateral root development | *A. thaliana* | 26163695 |
| IDA | LRP* | LRP* | Lateral root development | *A. thaliana* | 23479623 |
| JEKYLL | Flowers<br>Nucellus | Nucellus | Embryo/endosperm development | *H. vulgare* | 16766690 |
| KOD | Embryos | Embryo suspensor | Embryo suspensor development | *A. thaliana* | 26195729 |
| MEG1 | Endosperm transfer cells | Endosperm transfer cells | Transfer cell development | *Z. mays* | 26195729 |
| PEP7 | Roots | Roots | Lateral root development | *A. thaliana* | 36131543 |
| PSK,<br>PSY1 | Roots | Roots | Root elongation | *A. thaliana* | 20798316 |
| RGF1 | RAM* | RAM* | RAM* development | *A. thaliana* | 20798316 |
| RALF | Roots<br>Petioles | Roots | Root development | *N. attenuata*<br>*A. thaliana* | 17916115<br>24458638 |
| RALF1 | Roots | Roots | Root nodule development | *M. truncatula* | 18616408 |
| RALF4/19 | Pollen tubes | Pollen tubes | Pollen tube development | *A. thaliana* | 29242234 |

\* ESR: Embryo-surrounding region; SAM: Shoot apical meristem; RAM: Root apical meristem; LRP: Lateral root primordia

**Table S2.** The number of the previously identified vascular sap peptides and precursors in *Populus trichocarpa* and *Eucalyptus grandis* under different selection criteria (Chen et al., 2024).

| Characteristics | *P. trichocarpa* | *E. grandis* |
| --- | --- | --- |
|  | Peptides / Precursors | Peptides / Precursors |
| Vascular sap | 1997 / 801 | 331 / 229 |
| Conserved in Ptr & Egr<sup>*</sup><br>(≥ 70% peptide sequence identity) | 246 / 133 | 111 / 96 |
| Identical in Ptr & Egr<sup>†</sup><br>(100% peptide sequence identity) | 28 / 31 | 28 / 27 |
| Identical & xylem-specific | 17 / 16 | 17 / 14 |

\* 70% sequence identity was determined based on the length of the shorter peptide between *P. trichocarpa* and *E. grandis*. Ptr and Egr represent *P. trichocarpa* and *E. grandis*, respectively. † Peptides with identical sequences and lengths in both *P. trichocarpa* and *E. grandis*.

## References

Almagro Armenteros JJ, Tsirigos KD, Sonderby CK, Petersen TN, Winther O, Brunak S, von Heijne G, Nielsen H (2019) SignalP 5.0 improves signal peptide predictions using deep neural networks. Nat. Biotechnol. 37: 420–423

Altschul SF, Madden TL, Schäffer AA, Zhang J, Zhang Z, Miller W, Lipman DJ (1997) Gapped BLAST and PSI-BLAST: a new generation of protein database search programs. Nucleic Acids Res. 25: 3389–3402

Alvarez S, Goodger JQ, Marsh EL, Chen S, Asirvatham VS, Schachtman DP (2006) Characterization of the maize xylem sap proteome. J. Proteome Res. 5: 963–972

Anne P, Amiguet-Vercher A, Brandt B, Kalmbach L, Geldner N, Hothorn M, Hardtke CS (2018) CLERK is a novel receptor kinase required for sensing of root-active CLE peptides in *Arabidopsis*. Development 145: dev162354

Bollhöner B, Jokipii-Lukkari S, Bygdell J, Stael S, Adriasola M, Muñiz L, Van Breusegem F, Ezcurra I, Wingsle G, Tuominen H (2018) The function of two type II metacaspases in woody tissues of *Populus* trees. New Phytol. 217: 1551–1565

Buono RA, Hudecek R, Nowack MK (2019) Plant proteases during developmental programmed cell death. J. Exp. Bot. 70: 2097–2112

Butenko MA, Patterson SE, Grini PE, Stenvik GE, Amundsen SS, Mandal A, Aalen RB (2003) Inflorescence deficient in abscission controls floral organ abscission in *Arabidopsis* and identifies a novel family of putative ligands in plants. Plant Cell 15: 2296–2307

Carella P, Wilson DC, Kempthorne CJ, Cameron RK (2016) Vascular sap proteomics: providing insight into long-distance signaling during stress. Front. Plant Sci. 7: 651

Chang S, Puryear J, Cairney J (1993) A simple and efficient method for isolating RNA from pine trees. Plant Mol. Biol. Rep. 11: 113–116

Chen CH, Liou PC, Hsu YF, Wang IF, Kuo CY, Huang KH, Yu JH, Chen CW, Wu CC, Lin DG, et al. (2024) A sap peptide conserved across flowering plants positively regulates lignin biosynthesis, biomass and immunity. BioRxiv: 2024.05.20.594799

Chen H, Wang JP, Liu H, Li H, Lin YCJ, Shi R, Yang C, Gao J, Zhou C, Li Q, et al. (2019) Hierarchical transcription factor and chromatin binding network for wood formation in *Populus trichocarpa*. Plant Cell 31: 602–626

Chen S, Zhou Y, Chen Y, Gu J (2018) fastp: an ultra-fast all-in-one FASTQ preprocessor. Bioinformatics 34: i884–i890

Chen YL, Hsieh JWA, Kuo SC, Kao CT, Tung CC, Yu JH, Chang TH, Ku C, Xie J, Zhang D, et al. (2024) Merit of integrating in situ transcriptomics and anatomical information for cell annotation and lineage construction in single-cell analyses of *Populus*. Genome Biol. 25: 85

Chen YL, Lee CY, Cheng KT, Chang WH, Huang RN, Nam HG, Chen YR (2014) Quantitative peptidomics study reveals that a wound-induced peptide from PR-1 regulates immune signaling in tomato. Plant Cell 26: 4135–4148

Chen YL, Lin FW, Cheng KT, Chang CH, Hung SC, Efferth T, Chen YR (2023) XCP1 cleaves Pathogenesis-related protein 1 into CAPE9 for systemic immunity in *Arabidopsis*. Nat. Commun. 14: 4697

Chen YL, Fan KT, Hung SC, Chen YR (2020) The role of peptides cleaved from protein precursors in eliciting plant stress reactions. New Phytol. 225: 2267–2282

Chung KP, Zeng Y (2017) An overview of protein secretion in plant cells. Methods Mol. Biol. 1662: 19–32

Coll NS, Vercammen D, Smidler A, Clover C, Van Breusegem F, Dangl JL, Epple P (2010) *Arabidopsis* type I metacaspases control cell death. Science 330: 1393–1397

Cronk QCB, Forest F (2017) “The evolution of angiosperm trees: from palaeobotany to genomics”, in Groover A, Cronk QCB (eds.) Comparative and evolutionary genomics of angiosperm trees. Switzerland: Springer, Cham, pp. 1–17

Crooks GE, Hon G, Chandonia JM, Brenner SE (2004) WebLogo: a sequence logo generator. Genome Res. 14: 1188–1190

Dafoe NJ, Constabel CP (2009) Proteomic analysis of hybrid poplar xylem sap. Phytochemistry 70: 856–863

De Smet I, Voss U, Jürgens G, Beeckman T (2009) Receptor-like kinases shape the plant. Nat. Cell Biol. 11: 1166–1173

Demichev V, Messner CB, Vernardis SI, Lilley KS, Ralser M (2020) DIA-NN: neural networks and interference correction enable deep proteome coverage in high throughput. Nat. Methods 17: 41–44

Djordjevic MA, Oakes M, Wong CE, Singh M, Bhalla P, Kusumawati L, Imin N (2011) Border sequences of *Medicago truncatula* CLE36 are specifically cleaved by endoproteases common to the extracellular fluids of *Medicago* and soybean. J. Exp. Bot. 62: 4649–4659

Elias JE, Gygi SP (2007) Target-decoy search strategy for increased confidence in large-scale protein identifications by mass spectrometry. Nat. Methods 4: 207–214

Etchells JP, Mishra LS, Kumar M, Campbell L, Turner SR (2015) Wood formation in trees is increased by manipulating PXY-regulated cell division. Curr. Biol. 25: 1050–1055

Etchells JP, Turner SR (2010) The PXY-CLE41 receptor ligand pair defines a multifunctional pathway that controls the rate and orientation of vascular cell division. Development 137: 767–774

Farrokhi N, Whitelegge JP, Brusslan JA (2008) Plant peptides and peptidomics. Plant Biotechnol. J. 6: 105–134

Fukuda H (2004) Signals that control plant vascular cell differentiation. Nat. Rev. Mol. Cell Biol. 5: 379–391

Fukuda H, Hardtke CS (2020) Peptide signaling pathways in vascular differentiation. Plant Physiol. 182: 1636–1644

Fukuda H, Ohashi-Ito K (2019) Vascular tissue development in plants. Curr. Top. Dev. Biol. 131: 141–160

Funk V, Kositsup B, Zhao C, Beers EP (2002) The *Arabidopsis* xylem peptidase XCP1 is a tracheary element vacuolar protein that may be a papain ortholog. Plant Physiol. 128: 84–94

Ge Z, Bergonci T, Zhao Y, Zou Y, Du S, Liu MC, Luo X, Ruan H, Garcia-Valencia LE, Zhong S, et al. (2017) *Arabidopsis* pollen tube integrity and sperm release are regulated by RALF-mediated signaling. Science 358: 1596–1600

Goad DM, Zhu C, Kellogg EA (2017) Comprehensive identification and clustering of CLV3/ESR-related (CLE) genes in plants finds groups with potentially shared function. New Phytol. 216: 605–616

Goodstein DM, Shu S, Howson R, Neupane R, Hayes RD, Fazo J, Mitros T, Dirks W, Hellsten U, Putnam N, et al. (2012) Phytozome: a comparative platform for green plant genomics. Nucleic Acids Res. 40: D1178–D1186

Hander T, Fernández-Fernández ÁD, Kumpf RP, Willems P, Schatowitz H, Rombaut D, Staes A, Nolf J, Pottie R, Yao P, et al. (2019) Damage on plants activates Ca^2+^-dependent metacaspases for release of immunomodulatory peptides. Science 363: eaar7486

Hazak O, Brandt B, Cattaneo P, Santiago J, Rodriguez-Villalon A, Hothorn M, Hardtke CS (2017) Perception of root-active CLE peptides requires CORYNE function in the phloem vasculature. EMBO Rep. 18: 1367–1381

Hirakawa Y, Sawa S (2019) Diverse function of plant peptide hormones in local signaling and development. Curr. Opin. Plant Biol. 51: 81–87

Hirakawa Y, Shinohara H, Kondo Y, Inoue A, Nakanomyo I, Ogawa M, Sawa S, Ohashi-Ito K, Matsubayashi Y, Fukuda H (2008) Non-cell-autonomous control of vascular stem cell fate by a CLE peptide/receptor system. Proc. Natl. Acad. Sci. U.S.A. 105: 15208–15213

Hou S, Liu D, He P (2021) Phytocytokines function as immunological modulators of plant immunity. Stress Biol. 1: 8

Hunt L, Bailey KJ, Gray JE (2010) The signalling peptide EPFL9 is a positive regulator of stomatal development. New Phytol. 186: 609–614

Ito Y, Nakanomyo I, Motose H, Iwamoto K, Sawa S, Dohmae N, Fukuda H (2006) Dodeca-CLE peptides as suppressors of plant stem cell differentiation. Science 313: 842–845

Katsir L, Davies KA, Bergmann DC, Laux T (2011) Peptide signaling in plant development. Curr. Biol. 21: R356–R364

Kim MJ, Jeon BW, Oh E, Seo PJ, Kim J (2021) Peptide signaling during plant reproduction. Trends Plant Sci. 26: 822–835

Kondo T, Sawa S, Kinoshita A, Mizuno S, Kakimoto T, Fukuda H, Sakagami Y (2006) A plant peptide encoded by *CLV3* identified by in situ MALDI-TOF MS analysis. Science 313: 845–848

Kubo M, Udagawa M, Nishikubo N, Horiguchi G, Yamaguchi M, Ito J, Mimura T, Fukuda H, Demura T (2005) Transcription switches for protoxylem and metaxylem vessel formation. Genes Dev. 19: 1855–1860

Li B, Dewey CN (2011) RSEM: accurate transcript quantification from RNA-Seq data with or without a reference genome. BMC Bioinform. 12: 323

Li Q, Lin YC, Sun YH, Song J, Chen H, Zhang XH, Sederoff RR, Chiang VL (2012) Splice variant of the SND1 transcription factor is a dominant negative of SND1 members and their regulation in *Populus trichocarpa*. Proc. Natl. Acad. Sci. U.S.A. 109: 14699–14704

Ligat L, Lauber E, Albenne C, San Clemente H, Valot B, Zivy M, Pont-Lezica R, Arlat M, Jamet E (2011) Analysis of the xylem sap proteome of *Brassica oleracea* reveals a high content in secreted proteins. Proteomics 11: 1798–1813

Lin YC, Li W, Sun YH, Kumari S, Wei H, Li Q, Tunlaya-Anukit S, Sederoff RR, Chiang VL (2013) SND1 transcription factor-directed quantitative functional hierarchical genetic regulatory network in wood formation in *Populus trichocarpa*. Plant Cell 25: 4324–4341

Lin YJ, Chen H, Li Q, Li W, Wang JP, Shi R, Tunlaya-Anukit S, Shuai P, Wang Z, Ma H, et al. (2017) Reciprocal cross-regulation of VND and SND multigene TF families for wood formation in *Populus trichocarpa*. Proc. Natl. Acad. Sci. U.S.A. 114: E9722–E9729

Love MI, Huber W, Anders S (2014) Moderated estimation of fold change and dispersion for RNA-seq data with DESeq2. Genome Biol. 15: 550

Lu KJ, van ’t Wout Hofland N, Mor E, Mutte S, Abrahams P, Kato H, Vandepoele K, Weijers D, De Rybel B (2020) Evolution of vascular plants through redeployment of ancient developmental regulators. Proc. Natl. Acad. Sci. U.S.A. 117: 733–740

Luo JS, Zhang Z (2019) Proteomic changes in the xylem sap of *Brassica napus* under cadmium stress and functional validation. BMC Plant Biol. 19: 280

Magallón S, Gómez-Acevedo S, Sánchez-Reyes LL, Hernández-Hernández T (2015) A metacalibrated time-tree documents the early rise of flowering plant phylogenetic diversity. New Phytol. 207: 437–453

Matsubayashi Y (2011) Post-translational modifications in secreted peptide hormones in plants. Plant Cell Physiol. 52: 5–13

Matsuzaki Y, Ogawa-Ohnishi M, Mori A, Matsubayashi Y (2010) Secreted peptide signals required for maintenance of root stem cell niche in *Arabidopsis*. Science 329: 1065–1067

Mitchum MG, Wang X, Davis EL (2008) Diverse and conserved roles of CLE peptides. Curr. Opin. Plant Biol. 11: 75–81

Mohd-Radzman NA, Laffont C, Ivanovici A, Patel N, Reid D, Stougaard J, Frugier F, Imin N, Djordjevic MA (2016) Different pathways act downstream of the CEP peptide receptor CRA2 to regulate lateral root and nodule development. Plant Physiol. 171: 2536–2548

Motomitsu A, Sawa S, Ishida T (2015) Plant peptide hormone signalling. Essays Biochem. 58: 115–131

Nakano Y, Yamaguchi M, Endo H, Rejab NA, Ohtani M (2015) NAC-MYB-based transcriptional regulation of secondary cell wall biosynthesis in land plants. Front. Plant Sci. 6: 288

Narvaez-Vasquez J, Ryan CA (2004) The cellular localization of prosystemin: a functional role for phloem parenchyma in systemic wound signaling. Planta 218: 360–369

Nemec-Venza Z, Madden C, Stewart A, Liu W, Novák O, Pěnčík A, Cuming AC, Kamisugi Y, Harrison CJ (2022) *CLAVATA* modulates auxin homeostasis and transport to regulate stem cell identity and plant shape in a moss. New Phytol. 234: 149–163

Ni J, Guo Y, Jin H, Hartsell J, Clark SE (2011) Characterization of a CLE processing activity. Plant Mol. Biol. 75: 67–75

Ohtani M, Akiyoshi N, Takenaka Y, Sano R, Demura T (2017) Evolution of plant conducting cells: perspectives from key regulators of vascular cell differentiation. J. Exp. Bot. 68: 17–26

Okamoto S, Suzuki T, Kawaguchi M, Higashiyama T, Matsubayashi Y (2015) A comprehensive strategy for identifying long-distance mobile peptides in xylem sap. Plant J. 84: 611–620

Okamoto S, Tabata R, Matsubayashi Y (2016) Long-distance peptide signaling essential for nutrient homeostasis in plants. Curr. Opin. Plant Biol. 34: 35–40

Okuda S, Suzuki T, Kanaoka MM, Mori H, Sasaki N, Higashiyama T (2013) Acquisition of LURE- binding activity at the pollen tube tip of *Torenia fournieri*. Mol. Plant 6: 1074–1090

Okuda S, Tsutsui H, Shiina K, Sprunck S, Takeuchi H, Yui R, Kasahara RD, Hamamura Y, Mizukami A, Susaki D, et al. (2009) Defensin-like polypeptide LUREs are pollen tube attractants secreted from synergid cells. Nature 458: 357–361

Pedrioli PGA (2010) “Trans-proteomic pipeline: a pipeline for proteomic analysis”, in Hubbard S, Jones A (eds.) Proteome Bioinformatics. Methods in Molecular Biology™ (Methods and Protocols). New Jersey: Humana Press, vol. 604, pp. 213–238

Perez-Riverol Y, Bai J, Bandla C, García-Seisdedos D, Hewapathirana S, Kamatchinathan S, Kundu DJ, Prakash A, Frericks-Zipper A, Eisenacher M, et al. (2022) The PRIDE database resources in 2022: a hub for mass spectrometry-based proteomics evidences. Nucleic Acids Res. 50: D543–D552

Pierre-Jerome E, Drapek C, Benfey PN (2018) Regulation of division and differentiation of plant stem cells. Annu. Rev. Cell Dev. Biol. 34: 289–310

Qian P, Song W, Yokoo T, Minobe A, Wang G, Ishida T, Sawa S, Chai J, Kakimoto T (2018) The CLE9/10 secretory peptide regulates stomatal and vascular development through distinct receptors. Nat. Plants 4: 1071–1081

Reece JB, Campbell NA (2011) Campbell Biology. Boston: Benjamin Cummings / Pearson

Ren SC, Song XF, Chen WQ, Lu R, Lucas WJ, Liu CM (2019) CLE25 peptide regulates phloem initiation in *Arabidopsis* through a CLERK-CLV2 receptor complex. J. Integr. Plant Biol. 61: 1043–1061

Ribeiro CL, Conde D, Balmant KM, Dervinis C, Johnson MG, McGrath AP, Szewczyk P, Unda F, Finegan CA, Schmidt HW, et al. (2020) The uncharacterized gene *EVE* contributes to vessel element dimensions in *Populus*. Proc. Natl. Acad. Sci. U.S.A. 117: 5059–5066

Sarkar AK, Luijten M, Miyashima S, Lenhard M, Hashimoto T, Nakajima K, Scheres B, Heidstra R, Laux T (2007) Conserved factors regulate signalling in *Arabidopsis thaliana* shoot and root stem cell organizers. Nature 446: 811–814

Sharma A, Hussain A, Mun BG, Imran QM, Falak N, Lee SU, Kim JY, Hong JK, Loake GJ, Ali A, et al. (2016) Comprehensive analysis of plant rapid alkalization factor (RALF) genes. Plant Physiol. Biochem. 106: 82–90

Shi R, Wang JP, Lin YC, Li Q, Sun YH, Chen H, Sederoff RR, Chiang VL (2017) Tissue and cell-type co-expression networks of transcription factors and wood component genes in *Populus trichocarpa*. Planta 245: 927–938

Stührwohldt N, Dahlke RI, Kutschmar A, Peng X, Sun MX, Sauter M (2015) Phytosulfokine peptide signaling controls pollen tube growth and funicular pollen tube guidance in *Arabidopsis thaliana*. Physiol. Plant. 153: 643–653

Strijk JS, Hinsinger DD, Zhang F, Cao K (2019) *Trochodendron aralioides*, the first chromosome-level draft genome in Trochodendrales and a valuable resource for basal eudicot research. GigaScience 8: giz136

Sugano SS, Shimada T, Imai Y, Okawa K, Tamai A, Mori M, Hara-Nishimura I (2010) Stomagen positively regulates stomatal density in *Arabidopsis*. Nature 463: 241–244

Takahashi F, Hanada K, Kondo T, Shinozaki K (2019) Hormone-like peptides and small coding genes in plant stress signaling and development. Curr. Opin. Plant Biol. 51: 88–95

Tameshige T, Ikematsu S, Torii KU, Uchida N (2017) Stem development through vascular tissues: EPFL-ERECTA family signaling that bounces in and out of phloem. J. Exp. Bot. 68: 45–53

Tang W, Kelley D, Ezcurra I, Cotter R, McCormick S (2004) LeSTIG1, an extracellular binding partner for the pollen receptor kinases LePRK1 and LePRK2, promotes pollen tube growth *in vitro*. Plant J. 39: 343–353

Tung CC, Kuo SC, Yang CL, Yu JH, Huang CE, Liou PC, Sun YH, Shuai P, Su JC, Ku C, et al. (2023) Single-cell transcriptomics unveils xylem cell development and evolution. Genome Biol. 24: 3

Uchida N, Lee JS, Horst RJ, Lai HH, Kajita R, Kakimoto T, Tasaka M, Torii KU (2012) Regulation of inflorescence architecture by intertissue layer ligand-receptor communication between endodermis and phloem. Proc. Natl. Acad. Sci. U.S.A. 109: 6337–6342

van de Meene AM, Doblin MS, Bacic A (2017) The plant secretory pathway seen through the lens of the cell wall. Protoplasma 254: 75–94

Von Groll U, Berger D, Altmann T (2002) The subtilisin-like serine protease SDD1 mediates cell-to-cell signaling during *Arabidopsis* stomatal development. Plant Cell 14: 1527–1539

Wang W, Vignani R, Scali M, Cresti M (2006) A universal and rapid protocol for protein extraction from recalcitrant plant tissues for proteomic analysis. Electrophoresis 27: 2782–2786

Wang Z, Mao Y, Guo Y, Gao J, Liu X, Li S, Lin YCJ, Chen H, Wang JP, Chiang VL, et al. (2020) MYB transcription factor161 mediates feedback regulation of *Secondary wall-associated NAC- Domain1* family genes for wood formation. Plant Physiol. 184: 1389–1406

Wrzaczek M, Vainonen JP, Stael S, Tsiatsiani L, Help-Rinta-Rahko H, Gauthier A, Kaufholdt D, Bollhöner B, Lamminmäki A, Staes A, et al. (2015) GRIM REAPER peptide binds to receptor kinase PRK5 to trigger cell death in *Arabidopsis*. EMBO J. 34: 55–66

Xu B, Ohtani M, Yamaguchi M, Toyooka K, Wakazaki M, Sato M, Kubo M, Nakano Y, Sano R, Hiwatashi Y, et al. (2014) Contribution of NAC transcription factors to plant adaptation to land. Science 343: 1505–1508

Yamaguchi M, Mitsuda N, Ohtani M, Ohme-Takagi M, Kato K, Demura T (2011) VASCULAR-RELATED NAC-DOMAIN7 directly regulates the expression of a broad range of genes for xylem vessel formation. Plant J. 66: 579–590

Yamaguchi M, Ohtani M, Mitsuda N, Kubo M, Ohme-Takagi M, Fukuda H, Demura T (2010) VND-INTERACTING2, a NAC domain transcription factor, negatively regulates xylem vessel formation in *Arabidopsis*. Plant Cell 22: 1249–1263

Yang JH, Lee KH, Du Q, Yang S, Yuan B, Qi L, Wang H (2020) A membrane-associated NAC domain transcription factor XVP interacts with TDIF co-receptor and regulates vascular meristem activity. New Phytol. 226: 59–74

Yeh CS, Wang Z, Miao F, Ma H, Kao CT, Hsu TS, Yu JH, Hung ET, Lin CC, Kuan CY, et al. (2019) A novel synthetic-genetic-array-based yeast one-hybrid system for high discovery rate and short processing time. Genome Res. 29: 1343–1351

Yu L, Di Q, Zhang D, Liu Y, Li X, Mysore KS, Wen J, Yan J, Luo L (2022) A legume-specific novel type of phytosulfokine, PSK-δ, promotes nodulation by enhancing nodule organogenesis. J. Exp. Bot. 73: 2698–2713

Yuan B, Wang H (2021) Peptide signaling pathways regulate plant vascular development. Front. Plant Sci. 12: 719606

Zhang H, Zhang H, Lin J (2020) Systemin-mediated long-distance systemic defense responses. New Phytol. 226: 1573–1582

Zhao Q, Gallego-Giraldo L, Wang H, Zeng Y, Ding SY, Chen F, Dixon RA (2010) An NAC transcription factor orchestrates multiple features of cell wall development in *Medicago truncatula*. Plant J. 63: 100–114

Zhong R, Demura T, Ye ZH (2006) SND1, a NAC domain transcription factor, is a key regulator of secondary wall synthesis in fibers of *Arabidopsis*. Plant Cell 18: 3158–3170

Zhong R, Lee C, Zhou J, McCarthy RL, Ye ZH (2008) A battery of transcription factors involved in the regulation of secondary cell wall biosynthesis in *Arabidopsis*. Plant Cell 20: 2763–2782

Zhu Y, Song D, Zhang R, Luo L, Cao S, Huang C, Sun J, Gui J, Li L (2020) A xylem-produced peptide PtrCLE20 inhibits vascular cambium activity in *Populus*. Plant Biotechnol. J. 18: 195–206

